# Retinal self-organization: a model of RGC and SAC mosaic formation

**DOI:** 10.1101/2021.10.22.465398

**Authors:** Jean de Montigny, Evelyne Sernagor, Roman Bauer

## Abstract

Individual retinal cell types exhibit semi-regular spatial patterns called retinal mosaics. These mosaics enable uniform sampling of visual information and are formed to varying degrees across cell types. Retinal ganglion cells (RGC) and amacrine cells (including starburst amacrine cells (SAC)) are notably known to exhibit such layouts. Mechanisms responsible for the formation of such organised structures and their requirements are still not well understood. Mosaic formation follows three main principles: (1) homotypic cells prevent nearby cells from adopting the same type, (2) cell tangential migration, with homotypic cell repulsion, (3) cell death (with RGCs exhibiting high rates of apoptosis).

Here, we use BioDynaMo, an agent-based simulation framework, to build a detailed and mechanistic model of mosaic formation. In particular, we investigate the implications of the three theories for RGC’s mosaic formation. We report that the cell migration mechanism yields the most regular mosaics and that cell death can create regular mosaics only if the death rate is kept below 30%, after which cell death has a negative impact on mosaic regularity. In addition, and in accordance with recent studies, we propose here that low density RGC type mosaics exhibit on average low regularities, and thus we question the relevance of regular spacing as a criterion for a group of RGCs to form a RGC type.

We also investigate SAC mosaics formation and possible interactions between the ganglion cell layer (GCL) and inner nuclear layer (INL) populations. Investigations are conducted both experimentally and by applying our simulation model to the SAC population. We report that homotypic interactions between the GCL and INL populations during mosaics creation are required to reproduce the observed SAC mosaics’ characteristics. This suggests that the GCL and INL populations of SACs might not be independent during retinal development.

**Author Summary:** Retinal function depends on cells self-organisation during early development. Understanding the mechanisms underlying this self-organisation could improve not only our comprehension of the retina and its development but also of the cortex. Ultimately, this could lead to novel therapeutic approaches for developmental diseases. Computational models can be of precious help to study this process of self-organisation, given that they are biologically plausible. In this sense, it is important that implemented developmental mechanisms follow the principle of locally available information, without any global knowledge or external supervisor. Here, we follow this principle to investigate mosaic formation during retinal development. In this work, we demonstrate that tangential migration is the only mechanism able to form regular mosaics and that the GCL/INL SAC populations might not be independent during their mosaic formation. More, we question the relevance of regular spacing for RGC types classification.

## Introduction

The mammalian retina is composed of six main types of neuronal cells (cones, rods, horizontal, bipolar, amacrine and ganglion cells), subdivided into many different anatomical and functional subtypes, forming a complexly organised structure. Notably, individual cell types exhibit semi-regular spatial patterns called retinal mosaics. Regular spacing between homotypic cells enable homogeneous processing of the light signals, leaving no perceptual holes within our visual field. In particular, sub-groups of retinal ganglion cells (RGCs) and star burst amacrine cells (SACs) are known to form regular mosaics and both cell types are widely used to study mosaic organization. SACs are divided into two populations, one located in the inner nuclear layer (INL), and the other in the ganglion cell layer (GCL). Each population forms an independent mosaic.

RGCs are located in the GCL and are the output cells of the retina, sending all visual information processed in the retina to the brain visual areas. There are many ways RGCs can be classified into subgroups. One simple, basic approach is to classify them into three functional and morphological groups depending on the sub-layer their dendrites laminate into in the inner plexiform layer (IPL), forming the On, Off and On-Off groups. On cells respond to increase of light while Off cells respond to a decrease of light and On-Off cells respond to both increase and decrease of light.

RGCs can however be divided into more than 40 types (1–3), each having different functional and anatomical characteristics. Their density is also known to greatsly differ, varying from less than 50 cells/mm^2^ to more than 300 cells/mm^2^ (3). It has been proposed that a group of RGCs has to fulfil four criteria in order to be considered a RGC type (3): 1. Morphological homogeneity (dendritic tree shape). 2. Identical physiological properties (electrophysiological response to light). 3. Similar gene expression (molecular signature). 4. Regular spacing (mosaic). Thus, being organised in mosaics could represent an important feature of each RGC type. Even if the total number of RGC types is estimated to more than 40, only 19 have been fully characterised (cellular density, morphology, molecular signature and functions) (3). Other RGC types have been only partially characterised.

RGCs are the first cell class to differentiate in the immature retina, generated in the ventricular zone, followed by migration to the GCL, where they start extending dendrites towards the IPL. RGCs are the only cell class notably more numerous in the immature retina than in the adult retina. Indeed, around 60% of newly born RGCs undergo programmed cell death (apoptosis) during the perinatal period (4). Interestingly, not much is known yet about the impact of RGC apoptosis on the maturation of retinal circuitry and visual pathways.

Despite being an important feature of retinal organization, retinal mosaic’ formation is not fully understood yet. In particular, three mechanisms are believed to potentially take part in their development: cell-fate determination (CF), programmed cell death (CD) and tangential cellular migration (CM) (2,5).

### Cell fate determination

CF is a process by which a cell of a certain type will prevent the emergence of same type cells in its immediate vicinity (6). After passing through an intrinsically determined state, retinal progenitors are still left in an undifferentiated state, but are now only capable of giving rise to a limited subset of cell types. The precise type the cells choose to differentiate into depends on extrinsic signals (7). These extrinsic signals can consist of chemical cues such as trans-membrane proteins (8–10) and may be delivered by an already differentiated retinal cell in order to block neighbouring undifferentiated cells from differentiating into the same cell type. This mechanism has been demonstrated in the retina (7) and is believed to be ubiquitous in the developing CNS.

### Programmed cell death

RGCs exhibit a very high rate of programmed death, or apoptosis (60-70% of the initial population (11)), during normal development. The CD mechanism is believed to be implicated in the selection of relevant cells in order to build a functional retina. Following this principle, cellular death has been proposed to be a consequence of RGCs not being able to establish correct axonal connections in the lateral geniculate nucleus in the thalamus (12). RGC cell death has also been shown to depend on neighbouring cells’ electrical activity (13,14). Creating either spatial or functional competition between homotypic cells could lead to the formation or refinement of mosaics. Due to major differences in death rate, the importance of programmed cell death upon mosaic formation seems however to vary between cell classes, and even between subgroups within the same cell class. Cell death has been proposed to contribute to mosaic formation for several cell groups in the retina, including amacrine cells (15,16) and at least one RGC type (14).

### Cellular migration

All retinal cells undergo migration during retinal development, both vertical (from one layer to another) and tangential (horizontal migration within the same layer). Cells can move between 20 and 100 μm tangentially from their initial location (17,18). This mechanism is believed to be implicated in mosaic formation. This has been demonstrated for SACs mosaic formation, where homotypic cells move tangentially away from each other (19). Tangential migration is believed to be a key mechanism in mosaic formation (15). Mechanisms responsible for this migration are not fully understood, even if chemical cues seem to play a key role, such as in the case of SACs (20). Diffusible signals or contact-mediated interactions between homotypic cells may be responsible for mosaic formation (21). It is important to note that mosaics appear, partially or completely, before extensive dendritic growth (21,22), and thus without contact-mediated interactions. However, other studies point out the importance of dendritic growth upon tangential migration (20,23,24). In all cases, cell-cell interactions seem to be mandatory for tangential migration.

Of course, it is likely that the formation of mosaic patterns is due to the combinations of all three mechanisms (22). Previous mathematical simulations of retinal mosaic formation have been conducted (5,13). These studies investigated the involvement of the CF, CD and CM mechanisms, suggesting a central role for the CM mechanism. However, these studies are highly abstract and do not mechanistically model retinal mosaic formation, thus limiting their biological relevance. No mechanistic or biologically plausible model of mosaic formation currently exists.

Agent-based (AB) simulation is a type of computational model in which each simulation object is an autonomous agent. Despite the absence of any global supervisor, highly complex structures can emerge from local interactions of agents that self-organise (25,26). It is a particularly relevant approach to model biological phenomena where cells exhibit this characteristic as well. Using the AB approach would allow the construction of mechanistic and realistic models of retinal mosaic formation.

The impact and implications of all mechanisms involved in mosaic formation (cell-fate, cell death, tangential migration) are not fully understood, and much remains to be done in order to establish the detailed mechanisms governing mosaic formation. In this work, we analyse mechanisms underlying retinal mosaics self-organisation using AB computational modelling. In particular, the biological requirements and the effect of individual mechanisms generating these cellular patterns are investigated.

## Results

### RGC mosaic development

We demonstrate here that a realistic AB implementation of the CF is able to significantly increase the mosaic regularity compared to a random distribution (p < 0.001). Indeed, as shown in Figure 1A, the average regularity index (RI, used to assess the regularity of the mosaics) values rapidly increase from random levels (between 1.8 and 2) until reaching a value of 2.42 (±0.09) at the end of the CF mechanism. However, such RI values are lower than the experimentally observed values (> 3), and so cannot be considered as solely responsible for the formation of regular mosaics.

**Figure 1:**
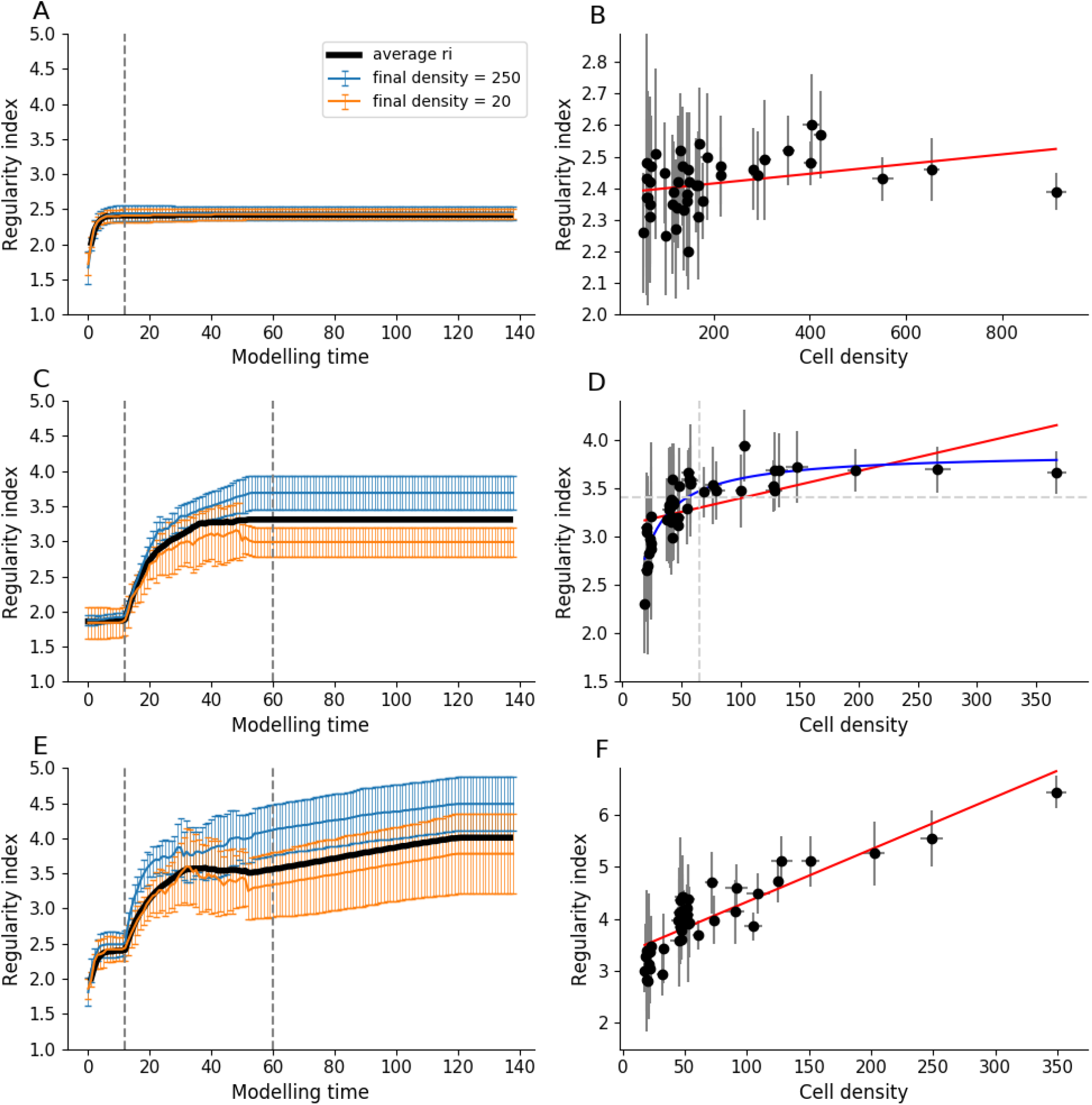
RGC mosaic formation modelling using an ABM approach. **A**,**C**,**E:** RI score evolution during simulation (x axis: 10 visualisation steps correspond to 1 developmental day in mouse). Average RI values for all RGC types are displayed in black while two populations of high and low densities (250 and 20 cells/mm 2 respectively) are displayed in blue and orange. The first vertical grey dashed line on the left indicates, if implemented, the end of the CF mechanism and if implemented, the beginning of the CD mechanism. The second grey dashed line indicates the end of the CD mechanism. **B**,**D**,**F:** Final RI score depending on cell density at the final step of the simulation. Error bars represent standard deviations for average RIs and densities. Red lines represent linear regressions (correlation coefficient: r=0.31, r=0.58 and r=0.87 for B, D and F respectively). The blue line in D represents a non-linear regression (a*x / b+x), while the horizontal dashed line represents the RI value under which no cell type of density higher than 125 is observed. **A**,**B:** CF mechanism only. **C**,**D:** CD mechanism only. **E**,**F:** Combination of CF, CD and CM combination.

Moreover, we find that the CF mechanism alone cannot explain high RI scores observed for some RGC types (> 5). As shown in Figure 1B, if CF is the only simulated mechanism, no correlation can be established between cell density and final RI values (correlation magnitude of 0.31). Thus, mosaics of high cell density reach similar RI values as seen in mosaics of low cell density, as illustrated by the blue and orange lines in Figure 1A.

The CD is also able to significantly increase RI compared to a random distribution (p<0.001), alone or in combination with the CF mechanism. As shown in Figure 1C, the average RI value increases from random (around 1.8) to 3.31 (±0.33) at the end of CD. This death rate amounts to around 65% when it reaches a steady state at the end of the simulation. These death rate dynamics are very similar to rates observed *in-vitro* (see Figure 2A). Moreover, and unlike for the case of the CF mechanism, CD is able to generate mosaics of medium regularities (RI > 3).

**Figure 2:**
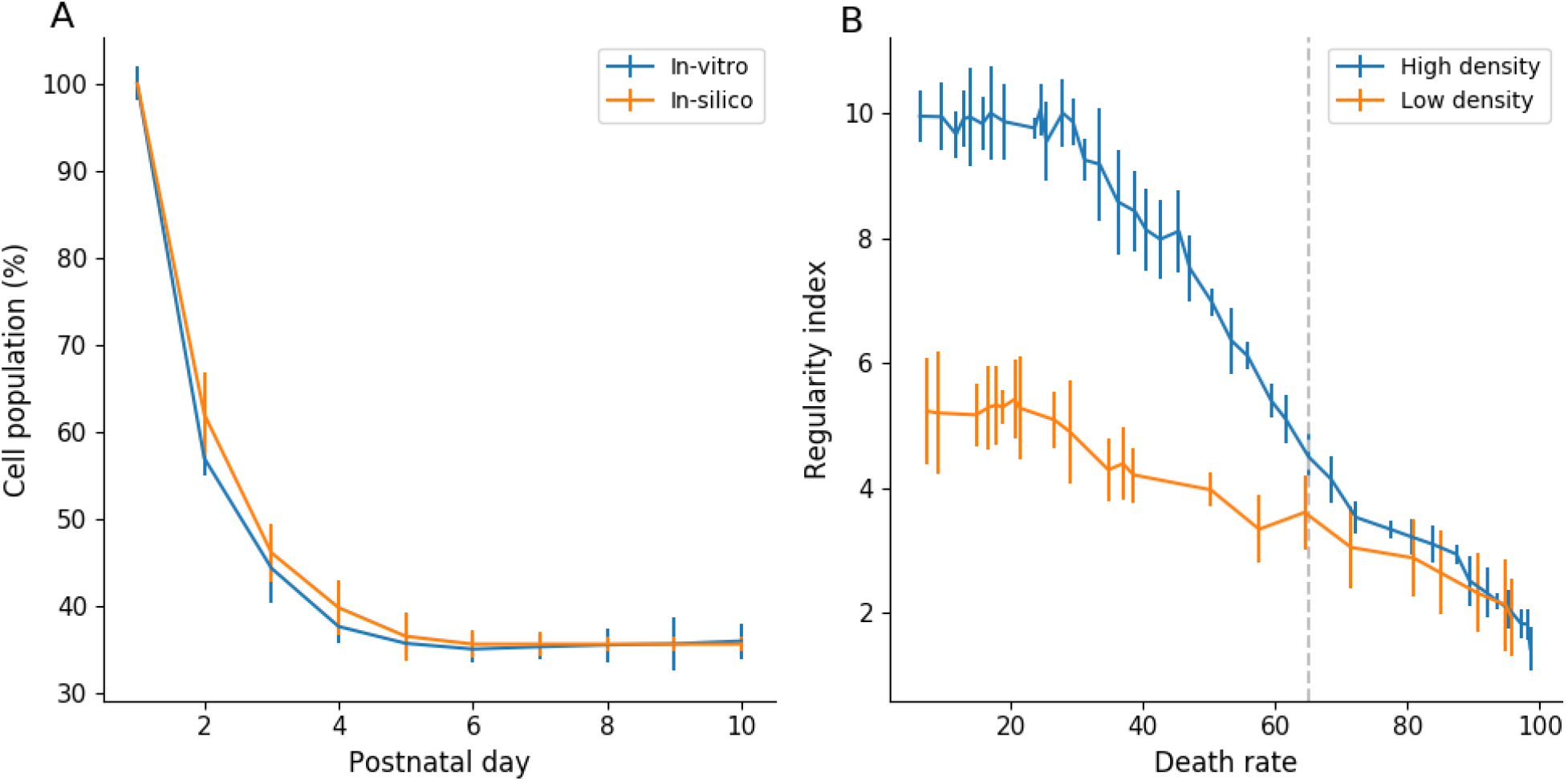
CD mechanism impact on RGC population. **A:** RGC population measured in-vitro (blue) and in simulations (orange). In-vitro population at day 1 is an estimation based on a final CD of 65%. Error bars represent standard deviation. **B:** RI score depending on final CD rate in a simulation implementing only the CD mechanism, for selected RGC populations of high density (blue curve, initial density = 571 cells/mm 2) and low density (orange curve, initial density = 114 cells/mm^2^). The vertical dotted line indicate a death rate of 65%. Error bars represent standard deviation.

Interestingly, and as shown by Figure 2B, the death rate measured *in-vitro* and selected for our simulations (grey vertical dashed line) is not the one generating the highest regularity. Indeed, the highest scores of RI are achieved for death rates between 5 and 30% of the RGC population, regardless of the initial density of the considered population. After 30% of cell death, RI decreases until it reaches a random distribution once around 90% of cell death is achieved. This is observed under both high and low-density conditions. This is observed in simulations with CD alone or in combination with CF. Interestingly, and in contrast to the CF mechanism, we observe strong differences between populations of high and low initial densities. As shown in Figure 1B, the high-density population is able to generate more regular mosaics than the population of low density, both for their maximum value (RI > 9 and RI > 5, respectively, when cell death is below 30%) and at 65% of cell death (RI = 4.49 ±0.36 and RI = 3.61 ±0.59, respectively). Therefore, a positive correlation between cell density and the final regularity is observed when the death rate is set to 65%. Mosaics of low density exhibit low RI values, while those of cell density higher than 65 cells/mm^2^ (vertical dashed line of Figure 1D) exhibit a higher average RI score of 3.35 (horizontal dashed line of Figure 1D).

While no differences are observed in RI scores between simulation of the CD mechanism and a combination of the CF and CD mechanisms if all mosaics are considered (3.31 ±0.33 and 3.48 ±0.44 respectively, p = 0.76), a positive impact on dense mosaics’ regularity (for cell densities higher than 125 cells/mm^2^) is to be noted. Thereby, RI values in the case of CF and CD combination plateaus around 4.1 instead of 3.6 if CD is the only implemented mechanism.

A combination of all three mechanisms (CF, CD and CM) is also able to generate mosaics significantly more regular than random distributions (p<0.001). Different steps of a simulation are illustrated by Figure 3.

**Figure 3:**
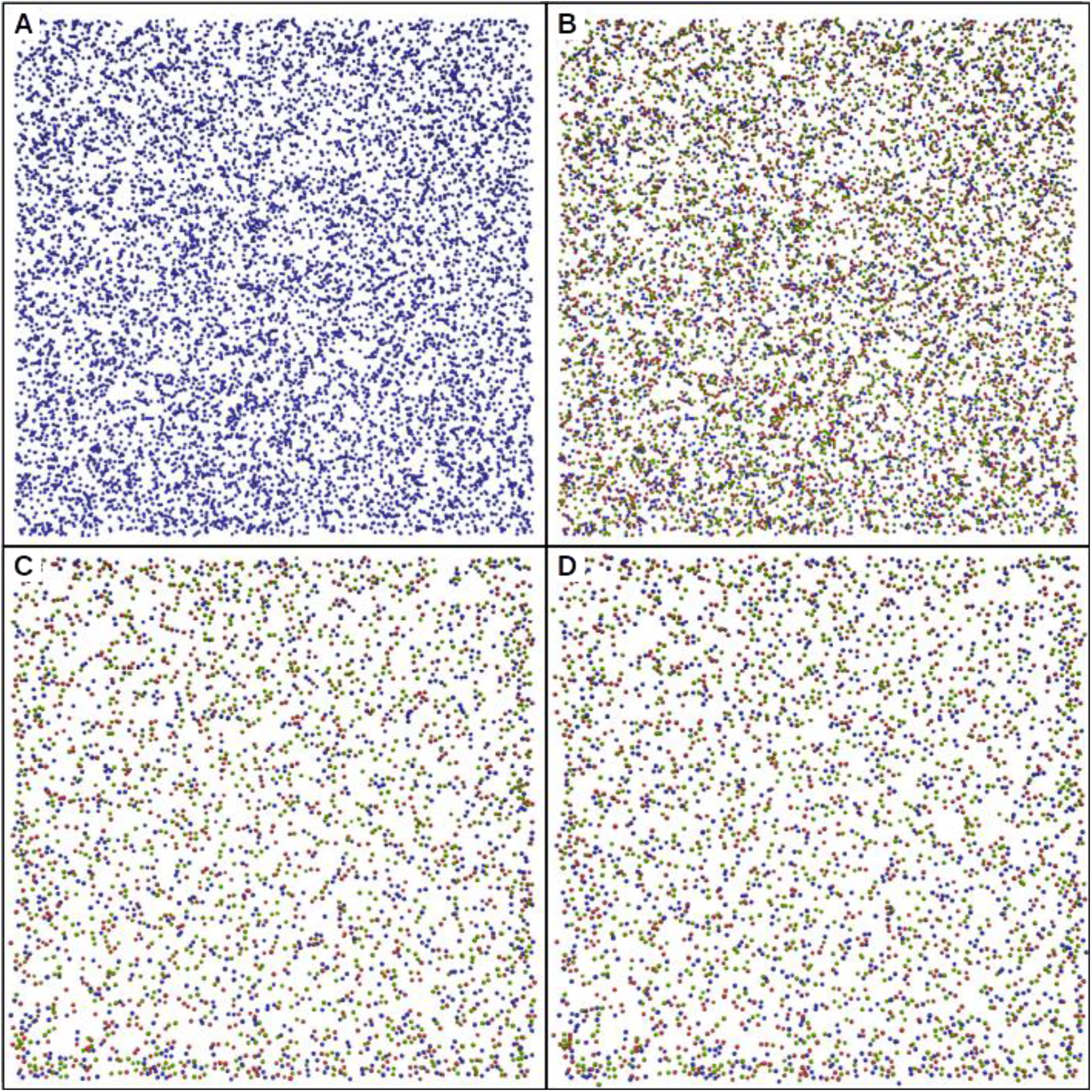
Time lapse of mosaic formation, using a combination of CF, CD and CM mechanisms using BioDynaMo. **A:** Simulation at step 0. All cells are undifferentiated and represented in blue. **B**,**C**,**D:** On cells are represented in green, Off cells in red and On-Off cells in blue. **B:** Simulation after the end of CF mechanism, at step 180. Average RI = 2.41. **C:** Simulation at the end of CD mechanism, step 1000. Average RI = 3.42. **D:** Simulation at the end of CM mechanism, step 2240. Average RI = 3.99.

As the CD and CM mechanisms require cells to be differentiated, CF is simulated beforehand. A first RI increase corresponding to the effect of CF is observed (see Figure 1E). After the CD and CM mechanisms are triggered (first dashed line), they give rise to a significant second increase, until the RI value stagnates toward the end of CD (simulation day 4 to 5.5 depending on the cell type). Finally, a third RI increase is observed after CD is over (second dashed line) due to the CM mechanism, leading to an average RI score of 4.01 (±0.75) at the end of the simulation. Unlike any other mechanism alone, and thanks to tangential migration, this simulation condition is able to generate highly regular mosaics (RI > 5). Moreover, a strong correlation appears between cell density and RI values (linear correlation magnitude of 0.87) as shown by Figure 1F. Thereby, only RGC types exhibiting a cell density higher than 125 cell/mm^2^ are able to generate mosaics with a RI value higher than 5. Thus, as illustrated by the blue and orange lines in Figure 1E, significant differences emerge between mosaics of high and low density. No significant differences are seen between simulations of CD and CM combination and simulations of CF, CD and CM combination.

When all three mechanisms are implemented, surviving cells migrate tangentially with an average distance of 8.72 μm (±0.11, n = 8 simulations), which is in accordance with *in-vivo* measurements reporting migration distance below 30µm (22). Important disparities in migration distance between cells are to be noted, as shown in Figure 4A, with an average migration distance standard deviation of 9.44 (±0.18). No correlation between final RI and migration distance can be seen. Likewise, no correlation appears between final density and migration distance if the whole population is considered. However, if only populations with a final density higher than 100 cells/mm 2 are considered, a strong correlation can be observed (correlation coefficient r = 0.92, see Figure 4B red line). Hence, the denser the cell type the larger the distance cells migrate.

**Figure 4:**
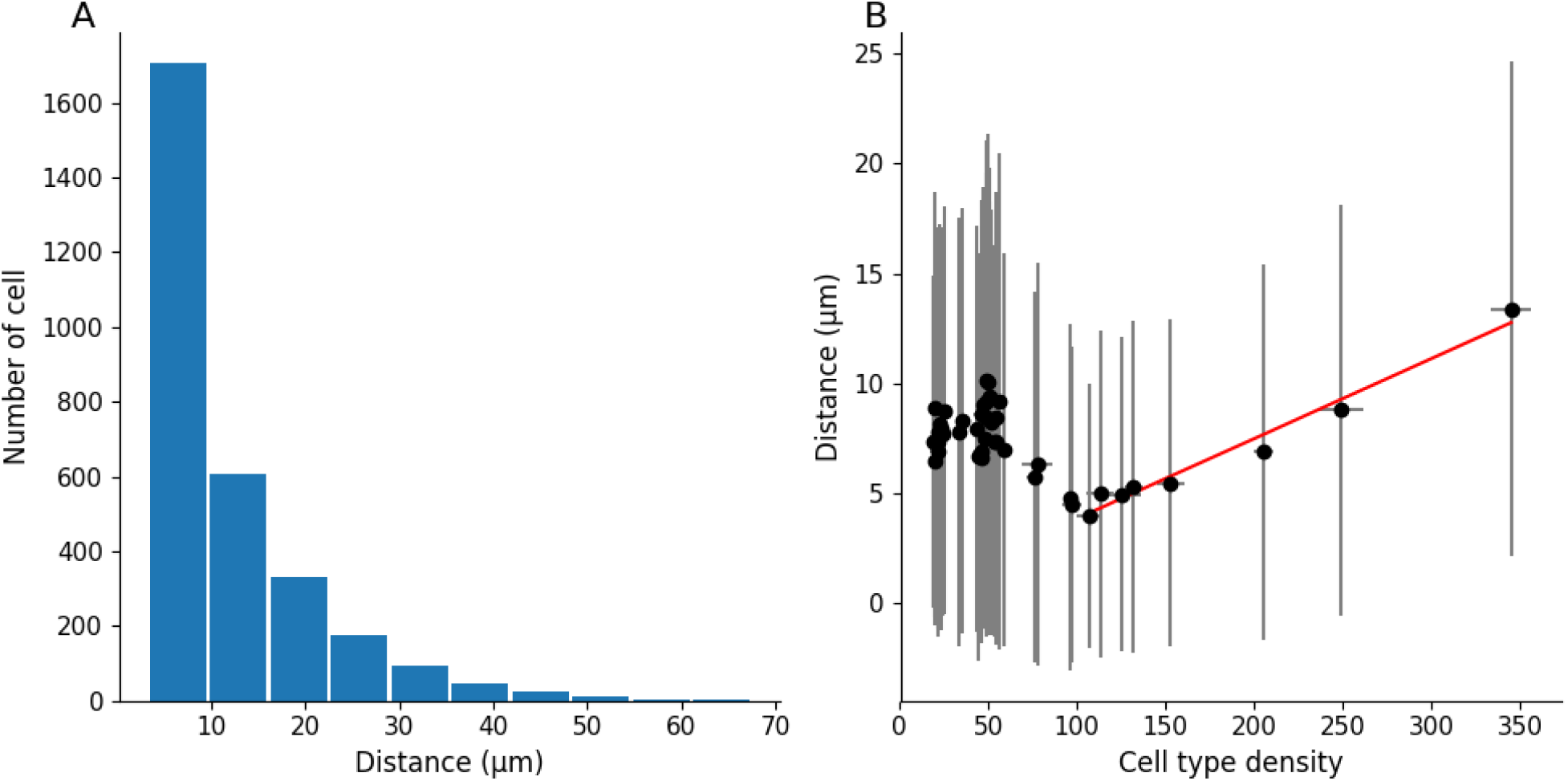
Migration distance measured in simulations implementing CF, CD and CM. **A:** Migration distance distribution. **B:** Relation between cell type density and migrating distance. The red line represents the correlation between migration distance and cell type density for densities higher than 100 cells/mm 2 (correlation coefficient r=0.92).

### SAC mosaic development

The SAC population is divided between two different cellular layers, the GCL and the INL, forming two separate populations (see Figure 5 for an illustration). Our *In-vitro* results exhibit no significant differences in the GCL and INL populations densities from P4 to P10 (p=0.27 and p=0.32 respectively, see Figure 6A). These two SAC populations exhibit regular pattern organisation, and no significant difference over time of their RI is measured from P4 to P10, as shown by Figure 6B. GCL and INL SAC mosaics are reported to be independent, in line with experimental data showing that SAC populations in the GCL and the INL only moderately overlap (20,27,28). A measure of these populations’ exclusion has then been conducted, showing no significant difference from P4 to P10, as shown by Figure 6C. This indicates that the INL and GCL SAC populations have already formed regular structures from P4 (shortly after GCL and INL separation), and do not exhibit further significant cell migration. This could indicate that mosaics are already formed and do not improve their regularity once SACs have migrated to their respective cellular layer. For this reason, SAC mosaics formation is implemented before GCL/INL separation in our simulations.

**Figure 5:**
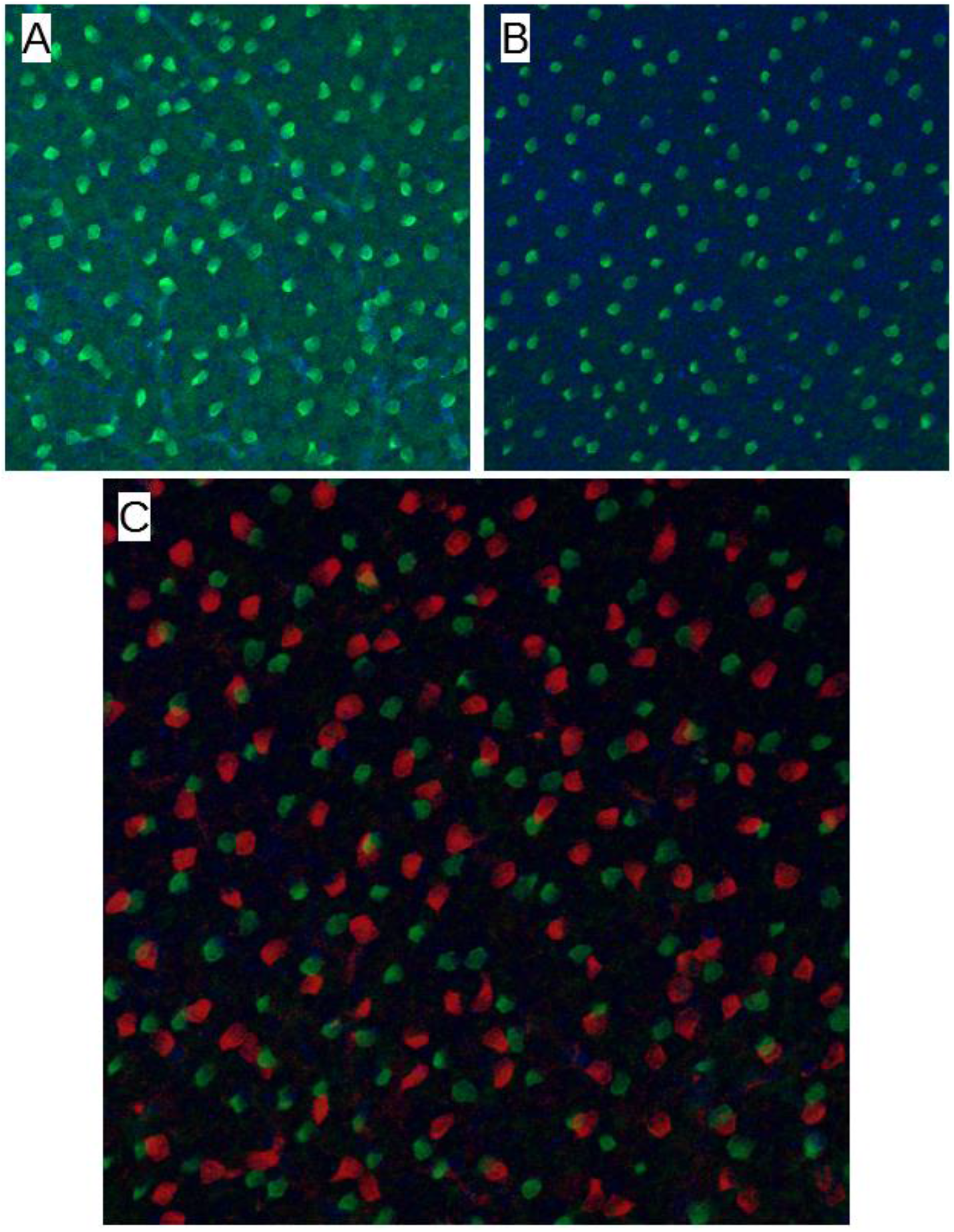
ChAT immunostaining on a P9 pup retina. **A:** GCL level. **B:** INL level. **C:** overlap of GCL (red) and INL (green) levels. GCL and INL level images are taken at the same x,y position, but at different depth focus. Regular SACs positioning can be observed in each cellular layer. Few cells overlap between GCL and INL levels are noted.

**Figure 6:**
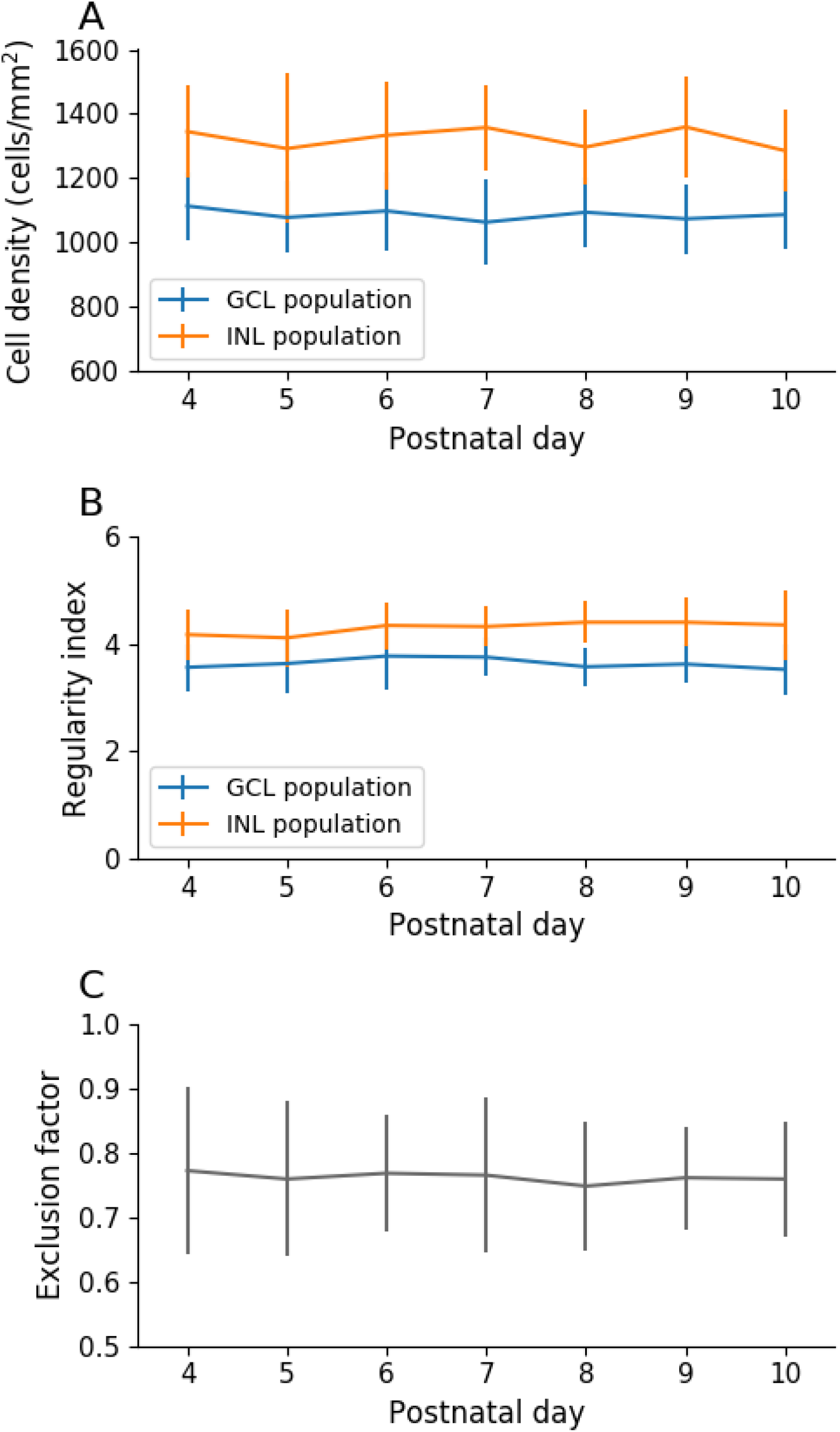
In-vitro SAC population characteristics through development. **A**,**B:** SACs in the CGL population are represented in blue, and the INL population is represented in orange. **A:** Cell density over time. **B:** Regularity index over time. **C: GCL and INL SAC population exclusion**. A score of 1 denotes two mosaics with a perfect exclusion and a score of 0 a total overlap of mosaics. Exclusion diameter of 32μm. Error bars represent standard deviation. P4: n=14; P5: n=15; P6: n=14; P7: n=12; P8: n=10; P9: n=9; P10: n=10. No differences are to be noted between P4 and P10.

Interestingly, by using an identical concentration threshold triggering CM for SAC in the GCL and INL, the GCL SAC population exhibits less regular mosaics than the INL population at the end of the simulation. This is observed in both developmental conditions, using either one common or two separate (one for the GCL population, one for the INL population) chemical substances for mosaic formation (RI of 3.57±0.12 and 4.11±0.12 respectively when one substance is used, RI of 3.36±0.07 and 4.37±0.15 respectively when two substances are used, n=8 for each group, p<0.0001). This mosaic regularity disparity is in accordance with observations in mouse (see Figure 6B) where the INL population has been reported to be more regular than in the GCL population. In our simulations, this disparity can be explained by the cell density difference between these two layers. Indeed, and as previously demonstrated in our simulations, the denser a cell population is, the more regular its mosaic can be. Hence, our model provides a mechanistic explanation for this observed difference in RIs between the two SAC populations.

However, we find that an important difference emerges between the two conditions concerning the exclusion factor of the two SACs populations: if one common developmental cue is used, GCL and INL mosaics exclude each other with a calculated exclusion factor of 0.71 (±0.01, n=8), similar to what has been measured *in-vitro* (0.74±0.09, n=5, see Figure 6). This indicates that the GCL and INL populations’ mosaics tend not to overlap, and so are not fully independent of each other. However, if two distinct developmental cues are used, the exclusion factor is lower, at 0.31 (±0.1, n=8), denoting independent mosaics that tend to overlap. In this second condition, the measured exclusion factor is significantly lower than the one observed in mouse (p<0.0001 using a T-test for two independent samples, n=8 and 5 respectively). Thus, only the first condition is able to reproduce the results observed *in-vitro*.

## Discussion

All computational simulations must be built upon biological data in order to offer relevant insight of a scientific problem. For this reason, information about retinal development has been gathered using in-vitro experimental observations. Thus, we followed RGCs characteristics through development using RBPMS staining in neonatal mouse retinas. This allowed us to measure death dynamic from P2 to P11. Using biological data from ours in-vitro experiments and from the literature, we built realistic simulations of retinal cells self-organization. This includes the number of RGCs types incorporated in our simulations, based on evidence from the literature (1–3). Notably, Sanes and Masland (2015) speculated that known RGC types represent only about 60% of the total RGC population, corresponding to around 1740 cells/mm^2^ from the total 3000 cells/mm^2^ observed in the mouse retina. In addition, it is important to note that from these known RGC populations, only 12.4% are On type. As On, Off and On-Off are equally numerous (30% to 35% each), a great number of On cells still needs to be discovered in order to reach the theoretical percentage of On RGC in the total RGC population. Thus, we can hypothesize that either: 1. Several high density On types have not yet been discovered. 2. There are more On types than Off or On-Off types.

The first hypothesis appears unlikely as RGCs are widely studied, especially with the emergence of large-scale and high density MEA recordings, but also using morphological and molecular characterisations. Thus, it is unlikely that the existence of dense On RGC types (representing the majority of the On population, and so being the most common On type) has not been captured by at least one of these techniques. The second hypothesis appears to be supported by experimental evidences because mice, similarly to other nocturnal animals, have rod-dominated vision. Indeed, rods are known to project their dendrites and to establish synaptic connections only to On bipolar cells, that in turn establish synaptic connections to On RGCs. In order to extract as many features as possible from a visual scene using mainly rod vision, a great diversity of specialized RGCs can be justified. The hypothesis of a great diversity of low density On types is also in agreement with Masland et al., (2015), who speculate that around 30 low density RGC types exist and are yet to be discovered. Baden et al. (2016) also estimate the total number of RGC types to be over 40, supporting the hypothesis of numerous low density RGC types, including On types. As it is still possible that On types of mid density has not been discovered, we chose to allow the possibility for this hypothesis in our simulations, in addition to adding multiple low density On RGCs.

One major basis of our simulations is the presence of chemical cues supporting cellular self-organization mechanisms. Evidences of such chemical cues have been previously reported (20,29,30).

### RGC mosaic formation

CF implication on RGC mosaics’ regularity is particularly difficult to study *in-vitro* or *in-vivo* as RGC progenitor cells do not express RGC type-specific markers cells will differentiate into. Despite experimental studies on RGC progenitors, no evidence has been found for RGC type-specific progenitors (31). Hence, RGC types are probably not pre-determined early on and so are likely to depend on extrinsic factors, such as the presence of chemical cues (7). Thereby, it allows for the contribution of a mechanism such as CF for RGC type differentiation, and its potential implication in mosaic formation. One major conclusion from our simulations is that highly regular mosaics (RI > 2.5) cannot be explained only through the CF mechanism. Likewise, the CF mechanism does not significantly increase the power of other mechanisms (CD and/or CM) neither. This suggests that RGC types are unlikely to be defined by cell body mosaics. They may instead be dictated by intrinsic factors (that still remain to be discovered), functional determination (dictated by the input from other cells), or a combination of intrinsic factors interacting with extrinsic factors.

In our simulations, the CD mechanism (alone, or in combination with the CF mechanism) is able to create regular mosaics (RI > 3.5) with a death rate of 65%. As this mechanism is based on a locally diffused chemical substance, homotypic cellular spacing (and therefore cell type initial density) has an important impact on the CD mechanism. For this reason, only populations with a high initial cell density exhibit regular mosaics. Importantly, our CD implementation is able to match measured RGC death dynamics during development, thus strengthening its plausibility. It should be pointed out that CD serves additional purposes in the retinal maturation process and is not only geared towards mosaic creation. Indeed, some cell types which do exhibit mosaic regularity do not undergo any significant levels of CD (such as horizontal cells or photoreceptors). In addition, as demonstrated here, the maximum positive impact of CD upon RI is reached at death rate lower than 30%, which is below the 60-70% death rate observed in mouse. This implies that even if CD can be involved in mosaic formation at early stages, cell death at levels above 30% is likely to be driven by other mechanisms and for other purposes than mosaic regularity. For instance, CD could be implicated in refining retinal functional connectivity and activity. CD could also have evolutionary advantage with regard to generating an optimised neural architecture (32).

Finally, CM is the only mechanism able to explain the formation of highly regular mosaics (RI > 5). As for the CD mechanism, the efficacy of the CM mechanism is dependent on cell density as it is based on local interactions. The shorter homotypic cellular distances are, the more they can sense and repulse each other. Thereby, a strong correlation emerges between RGC type populations densities and the regularity of their mosaics. Therefore, we propose here that low density RGC type mosaics exhibit on average significantly lower regularities than high density RGC types mosaics. It would be very informative to experimentally verify this prediction. To this date, this question remains unanswered. This hypothesis is in accordance with recent studies showing that some low density RGCs do not exhibit regular spacing (33). Moreover, we question here the relevance of regular spacing as a criterion for a group of RGC to form a RGC type. Indeed, if all low density RGC types do not exhibit highly regular spacing as predicted here, this criterion does not discriminate RGC types.

We show here that high mosaic regularity can be achieved with limited migration distance (8.72μm ±0.11 in average, n = 8). This average migration distance is in accordance with *in-vivo* measurements, reporting that RGCs and SACs tangential migration does not exceed 30μm (22). However, the average migration distance measured in our simulations is notably lower than the average migration distance experimentally measured at around 20μm (19) and could be explained by the absence of retinal surface expansion implementation in our simulations. The CM mechanism implemented here is based only on local cues and short-distance interactions, and thereby follows the description of tangential dispersion in mouse, reported as a local, short-distance, phenomenon (21). Our results are consistent with previous studies showing that a tangential cell dispersion does not appear to be directly related to the cell time of birth, but rather to its cell type (21).

The cellular migration and RI dynamics resulting from the CM mechanism are in agreement with the literature, where it is reported that RI increases mostly between P1 and P5, with the spacing between cells still increasing after that period, until P10 (2). After reaching the correct cell layers, a slower and finer tangential positioning phase of RGC within the GCL has been reported (19,34). Cellular movement during this period has been described as random but important for exact cellular positioning (34). In accordance with our results and as stated by other studies (27), these highly varied movements are likely to be related to mosaic formation and refinement. Indeed, these movements appear random as the whole RGC population (On, Off, On-Off population) is considered, while RGC populations should themselves be divided into types in order to meaningfully investigate RGCs lateral migration. If it were possible to examine each type independently, our model suggests that these movements, reported as random, would appear as coherent, as illustrated by Figure 7.

**Figure 7:**
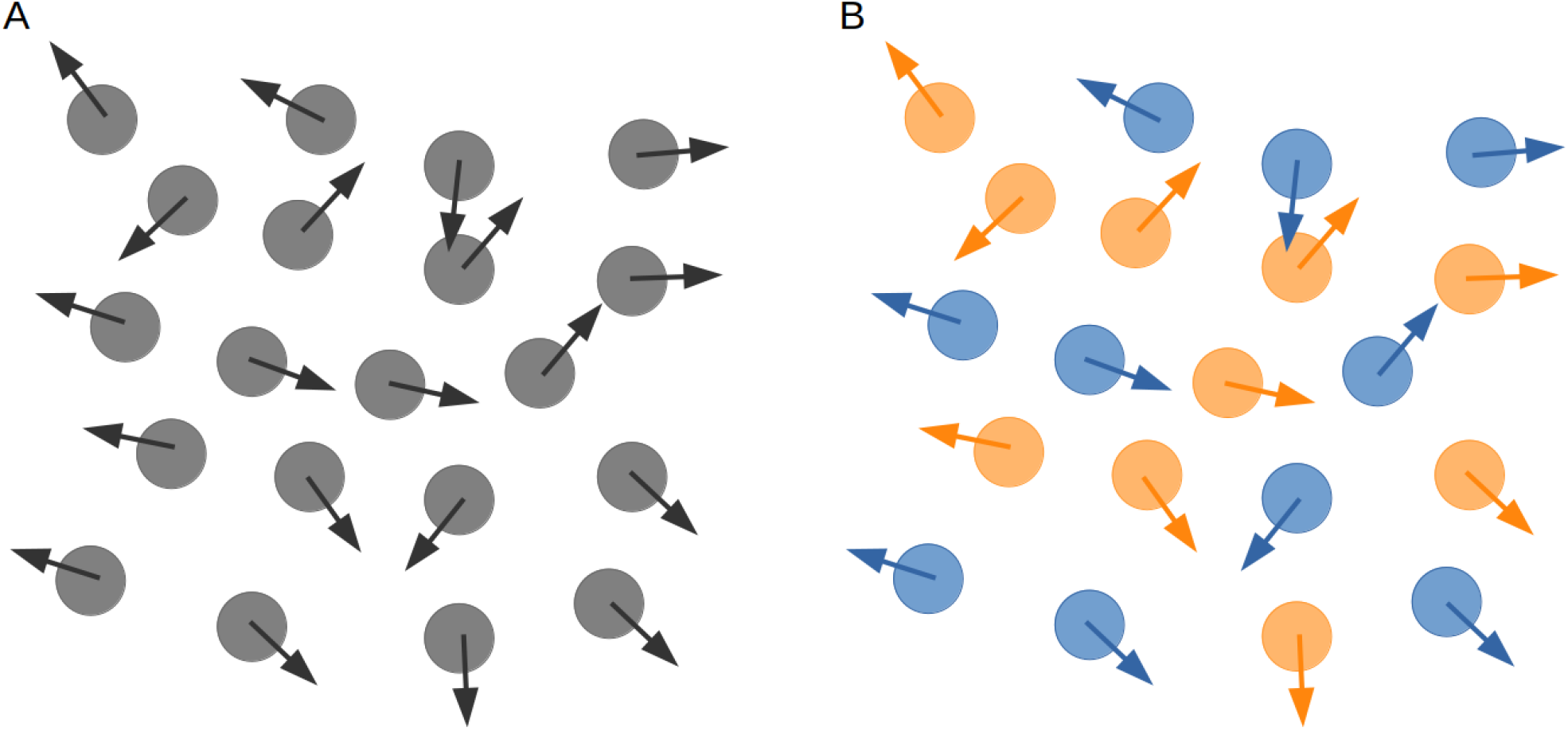
Cellular migration appearing as random if the whole population is considered homogeneously (**A**) or coherent (homotypic avoidance) if the population is sub-divided into two population (**B**).

### SAC mosaic formation

Beside most of RGCs, other retinal cell types are known to exhibit regular spacing, including SACs. This cell population is divided into the GCL and the INL. Both SAC layers form mosaics, that are reported to be independent from each other. Thus, there are only few overlaps between their populations. We found no RI variation from P3-4, indicating that these two SACs populations have already created their mosaics by P3-4, hence shortly after SACs migration into their respective cellular layer. In addition, we observed that the calculated exclusion factor does not vary, also supporting this assumption. Moreover, the observed complementarity of GCL and INL mosaics perhaps indicates interactions between these two SAC populations during their cellular organisation, before they migrate to their respective layer.

Here, we investigated this GCL/INL population interaction hypothesis further by building a simulation of SACs mosaics development. These simulations clearly show that our modelling procedure can successfully be applied to another cell population, without changing any simulation parameters. Indeed, we have been able to explain differences in GCL and INL mosaic regularities (the RI of the INL population being higher than that of the GCL population) by using only local interactions between SACs. This is the case if SACs constitute a unique population, or if GCL and INL populations are distinct (in other words, if one common or two distinct chemical cues are used). In the former case, this RI difference can be explained by the higher number of cells migrating to the INL compared to the GCL. Precisely, the percentage of a population characterised by a highly regular mosaic dictates the regularity of the resulting sub-population. Hence, the bigger the sub-population, the closer the obtained RI will be to the RI of the initial population, if cells constituting this sub-population are chosen randomly. For instance, if a population with a high RI is randomly divided into two sub-populations of 80% and 20% of the initial population (denoted respectively sub-population A and B), the sub-population A will have a RI closer to the initial population than the sub-population B. Thus, in the latter case, this observed RI difference between the GCL and INL populations can be explained by the higher cell density of SACs in the INL. This higher cell density in the INL allows more interactions and homotypic repulsion and thus the emergence of a higher RI than for the cells located in the GCL.

However, and importantly, only the simulation condition using a common chemical cue for mosaic formation is able to explain the complementarity observed between the GCL and INL populations. Indeed, if the two mosaics (GCL and INL) are formed independently, they largely overlap without exhibiting the mutual exclusion observed in-vitro. This suggests that the GCL and INL populations of SACs are not fully independent. Hence, our results predict that a shared guidance cue is responsible for mosaic formation of SACs in the GCL and INL. Locally diffused chemical (molecular) guidance could be a possible cue candidate for mosaic formation. If this is the case, our prediction could be potentially experimentally verified by using knock-out experiments blocking either the secretion or the reception of this chemical guidance.

## Methods

### Experimental work

#### Immunohistochemistry

Retinal wholemounts were prepared from mouse pups aged P2-P11, flattened on nitrocellulose membrane filters and fixed for 45 min in 4% paraformaldehyde. Retinas were then incubated in blocking solution — consisting in 5% of secondary antibody host species serum with 0.5% Triton X-100 in 0.1M phosphate buffer solution (PBS) — for 1 hour.

Retinas were incubated with 0.5% Triton X-100 with RBPMS (1:500) and ChAT (1:500) in PBS for 3 days at 4°C, then washed with PBS and incubated with 0.5% Triton X-100 with donkey anti rabbit Alexa 568 (1:500) and donkey anti goat Dylight 488 (1:500) in PBS for 1 day at 4°C. Finally, retinas were washed with PBS and embedded with OptiClear. Primary antibodies used were ChAT (AB144P, goat polyclonal, Merck Millipore) for SACs staining and RBPMS (1830-RBPMS, rabbit polyclonal, Phosphosolutions) for RGCs staining. Secondary antibodies used were Donkey anti rabbit Alexa 568 (A10042, Invitrogen) and Donkey anti goat Dylight 488 (SA5-10086, ThermoFisher Scientific).

Zeiss AxioImager with Apotome processing and the Zeiss LSM 800 confocal microscope were used to image the retinas. High-resolution of the whole retinal surface was achieved by imaging multiple individual adjacent areas. Individual images were subsequently stitched back together to view the entire retinal surface. Images at 40x magnification were also acquired in mid-peripheral regions in order to perform cell count and mosaic regularity measures.

#### Cell populations density

The average RGC and SAC density for each developmental day was measured by performing a manual cell count from P2 to P10 for RGCs and from P4 to P10 for SACs. By accounting for the surface expansion observed during retinal development, we estimated changes in populations through development. The estimated total RGC and SAC populations for a given retina are calculated by multiplying the averaged cell density (obtained from 3-6 sample areas per retina) by its corresponding retinal surface. These individual measurements are then averaged for each developmental day to give an estimation of the total population from P2 to P10. Cell population death rate during development is then calculated. In detail, the population of each retina measured on day D+1 is subtracted from the population of each retina measured on day D to calculate the amount of CD between day D and D+1. The amount of apoptosis measured between two consecutive days is then averaged to calculate the daily death rate of RGC and SAC populations. SAC populations in the GCL and INL are calculated separately.

#### SAC mosaics

Positions of SACs in the GCL and INL are also extracted in order to calculate mosaic regularities of these two populations from P4 to P10. A measure of GCL and INL mosaics exclusion has also been conducted. The calculated exclusion factor is based, for two distinct populations, on a count of cells from the first populations located within a determined distance (exclusion diameter) from cells belonging of the second population. This score is then normalised, to give an exclusion factor between 0 and 1. 1 denotes a perfect exclusion, meaning that all cells of the first population are located at a distance greater than the exclusion diameter from all cells of the second population. By consequence, only exclusion factors calculated with an identical exclusion diameter can be compared. A unique exclusion diameter of 32μm has been chosen here, corresponding to about 3 times the diameter of a SAC soma, and allowing a good discrimination between our different mosaics.

#### Ethics Statement

The experimental work was approved by the Animal Welfare Ethical Review Board (AWERB) of Newcastle University.

### BioDynaMo

Simulations were conducted using the agent-based simulation framework BioDynaMo (35). Each object in BioDynaMo is denoted as a simulation object, and possesses its own characteristics, such as its 3D geometry, mass, adherence and position in space. Individual neurons are represented by a sphere. Diffusion in 3D of chemical substances in the extracellular space has also been implemented, with the discrete central difference method. This diffusion is supported by grids representing substances concentration and gradients. Mechanical forces are also taken into account between all simulation objects such that they cannot overlap, but mechanically repulse each other. Each simulation object can have a *biology module* attached to it, that describes its behaviour at each simulation time step, such as substance secretion, cell migration or cell growth.

As an AB simulation framework, each simulation object is independent, without a central organisation unit that orchestrates the behaviour of all simulation objects. Thus, simulation objects only have access to their micro-environment, which consists of other simulation and chemical substances of the extracellular matrix in their proximity.

Several *biology modules* have been defined and used in our simulations, in order to describe cells behaviour for self-organisation (cell fate, cell death and cell migration) and chemical substances secretion.

### Simulations

All simulations took place in a cubic space of 1,300μm^3^, with cells of 7 to 8μm diameter randomly distributed (uniform distribution) in a space of 1,000μm×1,000μm×22μm. The initial cell density has been set to 8600 cells/mm^2^, in order to reach the RGC density once programmed CD mechanism is over — around 3000cells/mm^2^ reported in literature (3) and around 3500cells/mm^2^ in our measures. Mechanical interactions between simulation objects are taken into account, such that they cannot overlap, and mechanically repulse each other. The time step is set such that 160 steps simulate one day of development. Mosaic formation simulations run for a maximum of 2240 steps, corresponding to 14 days of development.

The global RGC population is subdivided into 43 types. Some have been precisely documented, such as the On or On-Off direction selective ganglion cells (DSGC), the local edge detector (LED), or the Off J-RGCs, and their population densities and dendritic arbours characteristics are known. However, these precisely documented RGC groups represent only 19 types, and merely about 60% of the total RGC population (∼1700 cells/mm^2^ over ∼3000cells/mm^2^) (3). RGC types composing the remaining 40% of the population have been estimated using results from Sanes and Masland (2015), Reese and Keeley (2015) and Baden et al. (2016). These authors state that numerous RGC types are still unknown and these cells are probably sparsely distributed across the retina. Thus, we implemented 24 additional RGC types of various but low densities. All implemented RGC types and their corresponding starting and final densities are summarised in Table 1.

**Table 1:** Implemented RGC types and parameters used for different conditions. **D:** death mechanism only. **FD:** fate and death mechanisms. **FDM:** fate, death and migration mechanisms. **Death:** concentration threshold for death mechanism. **Migration:** concentration threshold for migration mechanism. Parameters have been empirically chosen.

| Cell type | Type name | Start density<br>cells/mm<sup>2</sup> | Final density<br>cells/mm<sup>2</sup> | D<br>Death | FD<br>Death | FDM<br>Death | FDM<br>Migration |
| --- | --- | --- | --- | --- | --- | --- | --- |
| 0 | on-off_dsgca | 357 | 125 | 2.0367 | 2.0334 | 2.023 | 2.02 |
| 1 | on-off_dsgcb | 357 | 125 | 2.0367 | 2.0334 | 2.023 | 2.02 |
| 2 | on-off_dsgcc | 357 | 125 | 2.0367 | 2.0334 | 2.023 | 2.02 |
| 3 | on-off_dsgcd | 357 | 125 | 2.0367 | 2.0334 | 2.023 | 2.02 |
| 4 | on-off_m3 | 57 | 20 | 1.9872 | 1.9855 | 1.9855 | 1.983 |
| 5 | on-off_led | 714 | 250 | 2.116 | 2.098 | 2.08 | 2.065 |
| 6 | on-off_u | 57 | 20 | 1.9872 | 1.9855 | 1.985 | 1.983 |
| 7 | on-off_v | 57 | 20 | 1.9872 | 1.9855 | 1.985 | 1.983 |
| 8 | on-off_w | 171 | 60 | 2.001 | 1.9978 | 1.996 | 1.994 |
| 9 | on-off_x | 143 | 50 | 1.9968 | 1.9945 | 1.994 | 1.9925 |
| 10 | on-off_y | 114 | 40 | 1.994 | 1.993 | 1.993 | 1.991 |
| 11 | on-off_z | 114 | 40 | 1.994 | 1.993 | 1.993 | 1.991 |
| 100 | on_dsgca | 114 | 40 | 1.994 | 1.993 | 1.993 | 1.991 |
| 101 | on_dsgcb | 114 | 40 | 1.994 | 1.993 | 1.993 | 1.991 |
| 102 | on_dsgcc | 114 | 40 | 1.994 | 1.993 | 1.993 | 1.991 |
| 103 | on_aplha | 114 | 40 | 1.994 | 1.993 | 1.993 | 1.991 |
| 104 | on_m2 | 160 | 56 | 2 | 1.9953 | 1.994 | 1.992 |
| 105 | on_m4 | 57 | 20 | 1.9872 | 1.9855 | 1.985 | 1.983 |
| 106 | on_m5 | 57 | 20 | 1.9872 | 1.9855 | 1.985 | 1.983 |
| 107 | on_o | 428 | 150 | 2.05 | 2.0425 | 2.035 | 2.031 |
| 108 | on_p | 286 | 100 | 2.022 | 2.018 | 2.012 | 2.01 |
| 109 | on_q | 286 | 100 | 2.022 | 2.018 | 2.012 | 2.01 |
| 110 | on_r | 228 | 80 | 2.011 | 2.0082 | 2.004 | 2.002 |
| 111 | on_s | 171 | 60 | 2.001 | 1.9978 | 1.995 | 1.993 |
| 112 | on_t | 171 | 60 | 2.001 | 1.9978 | 1.995 | 1.993 |
| 113 | on_u | 143 | 50 | 1.9968 | 1.9945 | 1.994 | 1.9925 |
| 114 | on_v | 143 | 50 | 1.9968 | 1.9945 | 1.994 | 1.9925 |
| 115 | on_w | 97 | 34 | 1.993 | 1.9934 | 1.989 | 1.987 |
| 116 | on_x | 57 | 20 | 1.9872 | 1.9855 | 1.985 | 1.983 |
| 117 | on_y | 57 | 20 | 1.9872 | 1.9855 | 1.985 | 1.983 |
| 118 | on_z | 57 | 20 | 1.9872 | 1.9855 | 1.985 | 1.983 |
| 200 | off_aplhaa | 114 | 40 | 1.994 | 1.993 | 1.993 | 1.991 |
| 201 | off_aplhab | 114 | 40 | 1.994 | 1.993 | 1.993 | 1.991 |
| 202 | off_m1 | 180 | 63 | 2.006 | 1.9979 | 1.998 | 1.996 |
| 203 | off_j | 571 | 200 | 2.078 | 2.065 | 2.058 | 2.049 |
| 204 | off_mini_j | 1000 | 350 | 2.179 | 2.155 | 2.134 | 2.098 |
| 205 | off_midi_j | 228 | 80 | 2.011 | 2.0082 | 2.004 | 2.002 |
| 206 | off_u | 57 | 20 | 1.9872 | 1.9855 | 1.985 | 1.983 |
| 207 | off_v | 57 | 20 | 1.9872 | 1.9855 | 1.985 | 1.983 |
| 208 | off_w | 171 | 60 | 2.001 | 1.9978 | 1.995 | 1.993 |
| 209 | off_x | 143 | 50 | 1.9968 | 1.9945 | 1.994 | 1.9925 |
| 210 | off_y | 114 | 40 | 1.994 | 1.993 | 1.993 | 1.991 |
| 211 | off_z | 106 | 37 | 1.9935 | 1.9928 | 1.989 | 1.988 |

Cells are created with no predefined types when simulating the CF mechanism. Otherwise, cells of each RGC type are created matching their experimentally observed initial density.

#### Substance secretion

Each RGC type secretes a specific chemical substance that diffuses in the extracellular space, using grids of 2μm^3^ voxels. This secretion corresponds to an increase of substance concentration by 1 at the cell centre position. Undifferentiated cells do not secrete any substance.

#### RGC mosaic formation: CF

CF is implemented such that substances act as an inhibitor for cell differentiation, preventing nearby undifferentiated cells to adopt the same types. In this way, neighbouring cells are forced to differentiate into other RGC types. CF is the first event to occur during simulations, because CD and CM mechanisms operate on differentiated cells.

#### RGC mosaic formation: CD

The CD mechanism corresponds to the cells removing themselves from the simulation if their corresponding substance concentration is higher than a defined threshold. In this way, the clusters of homotypic cells exhibit high death rates and become sparser. As the cell density decreases, the initial multilayer collapses into a RGC monolayer. This is implemented as cells moving along the z axis toward the centre of the RGC layer, using their chemical cue. CD is triggered after completion of CF, and continues until a steady-state is reached, around a death rate of 65%. This steady-state is reached without global controllers but depends on the chosen concentration threshold triggering cell death. If this threshold is low the steady-state will be reached with a high death rate, while if this threshold is high the steady-state will be reached with a low death rate.

#### RGC mosaic formation: CM

CM is implemented such that the homotypic substances act as a repulsive factor. Thereby, cells exhibit short distance avoidance, moving tangentially against their substance gradient, distancing themselves from homotypic neighbours. We assume that CM is triggered after completion of CF, at the same time as CD, and continues either until a steady state or day 13 is reached.

Development conditions incorporating all combinations of these three mechanisms have been investigated. As the mechanisms influence each other, parameters vary depending on the implemented mechanisms. The CD mechanism parameters were chosen for each RGC type such that its final death rate is about 65%. The CM parameters were chosen depending on the CD parameter value and such that the interaction is kept to close range distance. Pseudocode corresponding to these three *biology modules* can be found in Pseudocode 1. Table 1 summarises the parameters used for RGC mosaics formation mechanisms.

**Pseudocode 1:**
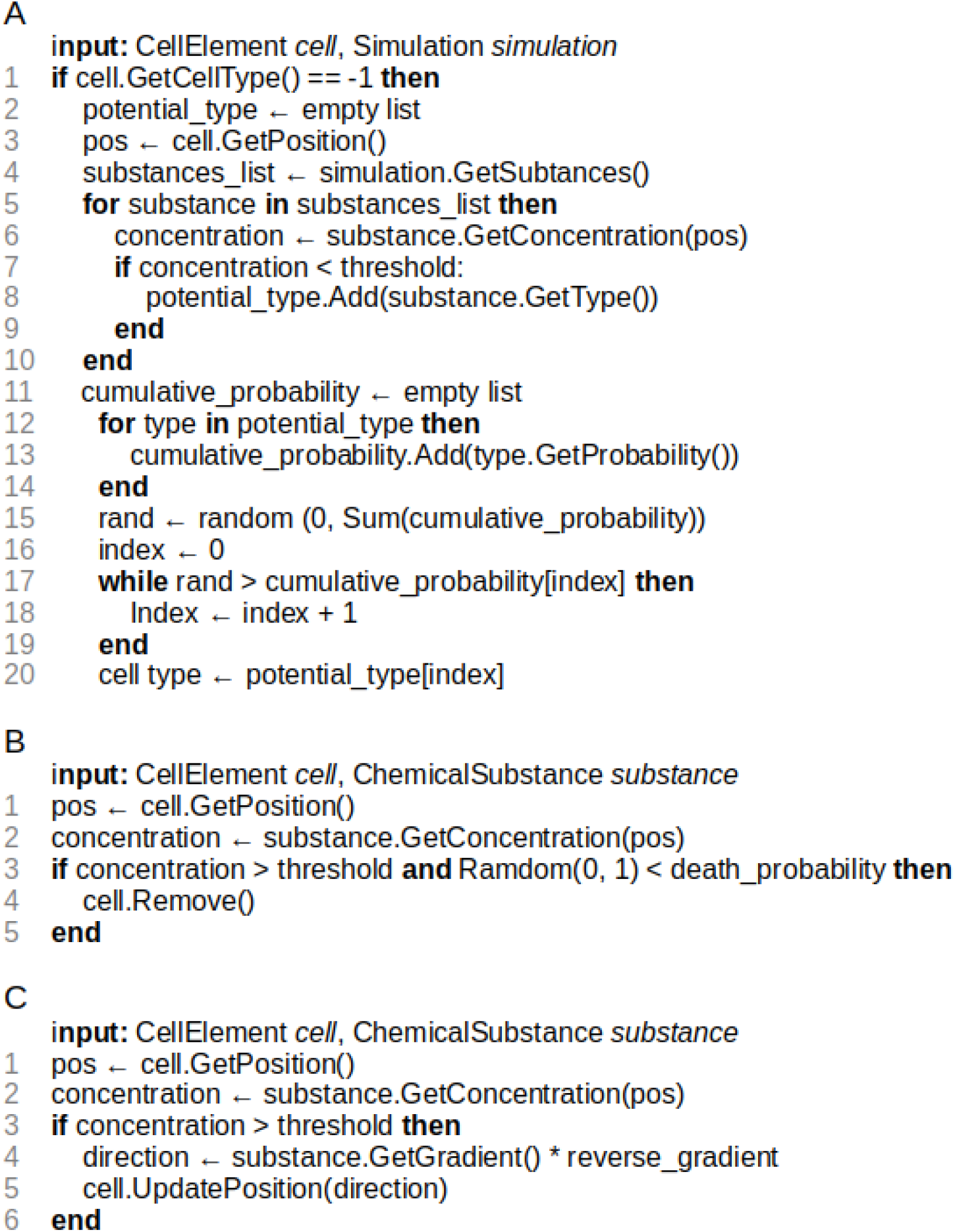
Pseudocode describing biology modules. **A:** cell fate. **B:** cell death. **C:** cell migration.

#### SAC mosaic formation

The simulation of SAC mosaic formation is achieved using locally diffused chemical cues triggering homotypic avoidance (tangential migration mechanism). Once mosaics are formed, the two populations migrate to their respective layers, along the Z axis. Two different developmental conditions have been implemented, using either one common or two separated (one for GCL population, one for INL population) chemical substances for mosaics formation. Importantly, concentration thresholds are identical for the GCL and INL populations. Parameters have been set such that the mosaic RIs match the measured RIs in mouse SACs mosaics.

### Data analysis

The RI was used to assess the regularity of the mosaics. It is computed as the average value of the closest neighbour distribution (distribution of the closest neighbour measured for each cell) divided by its standard deviation (36). The RI offers a single score that is able to discriminate regularity differences between mosaics of low regularities. In addition, and as previously reported by Reese and Keeley (2015), the RI offers a scale-invariant measure of mosaic regularity and thus more direct evidence of any change in the mosaic spatial organisation during development. It is not only the absolute RI value that carries information, but also its evolution across development, related to the contribution of each mosaic developmental mechanism (CF, CD, CM). However, it should be pointed out that RI is sensitive to a low sampling rate, leading to significant variability in RI scores for mosaics constituted of few cells.

Comparisons between two RI values have been conducted using T-tests for two independent samples.

## Author Contributions

JdM, ES and RB conceived and designed the experiments and wrote the paper. JdM performed the experiments and analyzed the data.

